# Harnessing *EGLN1* Gene Editing to Amplify HIF-1α and Enhance Human Angiogenic Response

**DOI:** 10.1101/2023.05.29.542734

**Authors:** Shahin Shams, Roberta S. Stilhano, Eduardo A. Silva

## Abstract

Therapeutic angiogenesis has been the focus of hundreds of clinical trials but approval for human treatment remains elusive. Current strategies often rely on the upregulation of a single proangiogenic factor, which fails to recapitulate the complex response needed in hypoxic tissues. Hypoxic oxygen tensions dramatically decrease the activity of hypoxia inducible factor prolyl hydroxylase 2 (PHD2), the primary oxygen sensing portion of the hypoxia inducible factor 1 alpha (HIF-1α) proangiogenic master regulatory pathway. Repressing PHD2 activity increases intracellular levels of HIF-1α and impacts the expression of hundreds of downstream genes directly associated with angiogenesis, cell survival, and tissue homeostasis. This study explores activating the HIF-1α pathway through *Sp*Cas9 knockout of the PHD2 encoding gene *EGLN1* as an innovative *in situ* therapeutic angiogenesis strategy for chronic vascular diseases. Our findings demonstrate that even low editing rates of *EGLN1* lead to a strong proangiogenic response regarding proangiogenic gene transcription, protein production, and protein secretion. In addition, we show that secreted factors of *EGLN1* edited cell cultures may enhance human endothelial cell neovascularization activity in the context of proliferation and motility. Altogether, this study reveals that *EGLN1* gene editing shows promise as a potential therapeutic angiogenesis strategy.

## INTRODUCTION

Therapeutic angiogenesis promotes new blood vessels via the delivery of pro-angiogenic factors and offers an alternative treatment for a wide range of ischemic cardiovascular diseases (CVDs) that conventional revascularization strategies have not addressed (1). Traditionally, therapeutic angiogenesis efforts focused on delivering one or more of either pharmacological compounds, growth factors, or proangiogenic cells to stimulate vascular network formation and restore blood flow to hypoxic tissues (2–6).

VEGF-A is a potent signaling protein that primarily targets endothelial cells, and is responsible for the proliferation and migration of these cells while mediating neovascular activity (7). VEGF-A has been extensively studied for therapeutic angiogenesis. Despite promising preclinical research, clinical studies looking at systemic administration of VEGF-A protein have shown disappointing clinical results and clinical gene therapy applications have likewise failed to reestablish blood flow in multiple ischemic vascular disease contexts (7,8). Moreover, long term overexpression of VEGF-A alone leads to unstable and immature vascular structures that can disrupt tissue homeostasis and exacerbate inflammation (9).

To overcome limitations related to the overexpression of a single factor such as VEGF-A, recent efforts have focused on pharmacologically and genetically manipulating the HIF-1α pathway, a master regulator of proangiogenic processes (10–15). The HIF-1α master regulatory pathway drives endogenous cellular hypoxic response, and activating it triggers modifications in the expression of hundreds of factors associated with angiogenesis and tissue homeostasis, leading to a coordinated hypoxic response for tissue organization, capillary formation, and blood vessel stabilization and maturation (16). Mechanistically, HIF-1α destabilization and degradation susceptibility is influenced by HIF prolyl hydroxylases (HIF-PHs), which requires oxygen for activity. When oxygen is depleted, HIF-PH suppression occurs, leading to HIF-1α protein stabilization (17). Stabilization of HIF-1α transcriptionally activates downstream genes that regulate glucose metabolism, cellular homeostasis, cell proliferation, cell migration, cell survival, ECM reorganization, and angiogenesis through the expression of more than a hundred growth factors, structural proteins, cytokines, and many other biologically active molecules relevant to therapeutic angiogenesis strategies (16,18).

Recent efforts to stabilize HIF-1α have been focused on pharmacological agents or gene delivery of degradation resistant HIF-1α variants (10–15). Specifically, the strategy of genetically silencing HIF-PHs has been explored, with HIF-PH 2 (PHD2), the most common variant encoded by the gene *EGLN1* (19,20). Global and conditional *EGLN1* knockout (KO) studies showed increased perfusion, capillary density, and resistance to CVD formation (21–25). In addition, human population studies of functional *EGLN1* mutations revealed improved responses to hypoxia and resulted in protection against some forms of CVD (26). In therapeutic angiogenesis contexts, *EGLN1* knockdown has shown great promise for various CVD models (19,20,27–29). However, no permanent strategies for *EGLN1* manipulation have been explored to establish a therapeutic angiogenesis intervention approach to date.

In this study, we explore if using *Sp*Cas9 to edit the *EGLN1* gene can permanently activate the HIF-1α pathway and trigger a robust downstream proangiogenic response in human cells (**Figure 1**). We hypothesized that long lasting *EGLN1* manipulations would lead to a robust increase in intracellular HIF-1α protein levels, yielding significant downstream effects – as measured by real time PCR for intracellular transcripts and ELISA for proangiogenic protein secretion. The experiments described in this study demonstrate that even marginal gene editing of *EGLN1* can induce a significant proangiogenic response. Both intracellular HIF-1α and secreted VEGF-A protein were significantly elevated in *EGLN1* KO populations, and the response of human endothelial cells to media preconditioned in *EGLN1* KO cultures were significantly increased compared to non-preconditioned controls. Taken together, this work, for the first time, indicates that permanent modification of *EGLN1* could be viable for developing an alternative therapeutic angiogenesis strategy that does not rely on traditional single factor overexpression.

**Figure 1.**
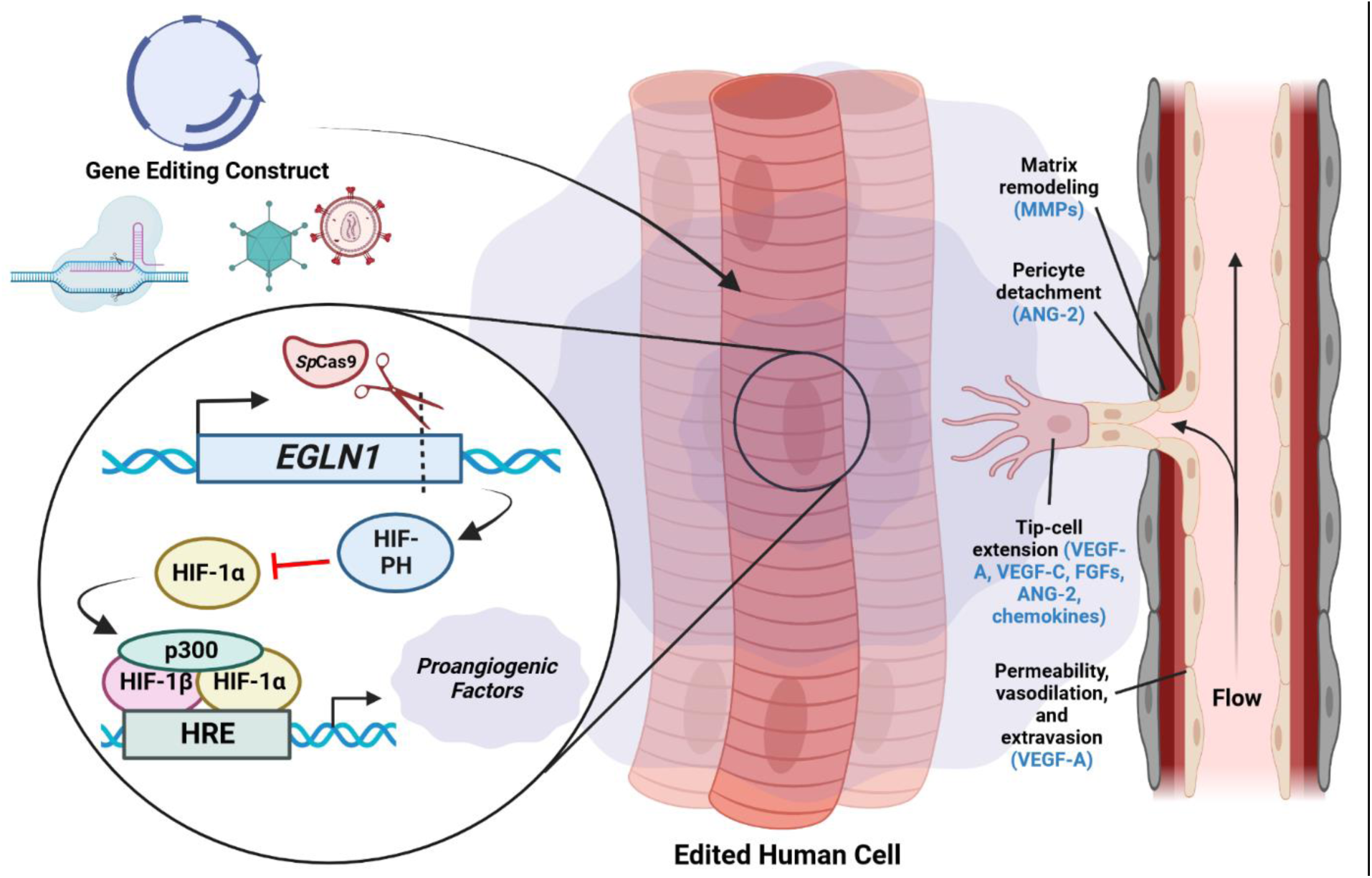
Gene editing activation of the HIF-1α proangiogenic cascade to initiate a neovascular response. Secreted proangiogenic factors regulated by HIF-1α include matrix metalloproteinases (MMPs), angiopoietin-2 (Ang2), vascular endothelial growth factors (VEGF-A and VEGF-C), fibroblast growth factors (FGFs), and chemokines.

## MATERIALS & METHODS

### *In silico* generation of human *EGLN1* sgRNA

*Sp*Cas9 sgRNA constructs localized to *EGLN1* were generated *in silico* via the CRISPR/Cas9 target site web tool CHOPCHOP (30). In brief, after entering the target gene, species of interest, chosen gene editing system, and desired mutation (such as knock-out, knock-in, activation, repression, etc.) into the web tool, the system identifies and ranks prospective guides. Consequently, the ranking is based on a multitude of factors, including predicted cleavage efficiency, self-complementarity, GC content, and predicted off-target effects.

### Production of *EGLN1* sgRNA expression cassettes

Lentiviral transfer vector plasmids containing *Sp*Cas9 guide scaffold sequences were selected as the mammalian expression cassettes of interest for both preliminary screening assays as well as VLP studies. Two plasmids were chosen for further development: a plasmid containing both an empty sgRNA scaffold as well as GFP (AddGene plasmid# 99375) with a final goal of VLP generation, and a plasmid containing both an empty sgRNA scaffold as well as a gene sequence encoding *Sp*Cas9 (AddGene plasmid# 57819) with a final goal of LV positive control vector generation. *In silico* generated guide sequences were cloned into both sgRNA lentiviral transfer vectors through site directed mutagenesis (New England Biolabs, Q5 Site Directed Mutagenesis Kit, Cat# E0554S). Proper insertion of the guide sequences was validated through Sanger Sequencing at the UC Davis Sequencing Core. Plasmids were produced in working concentrations in chemically competent *E. coli* and purified for cell culture application through commercially available kits (Qiagen, Maxi Plasmid Kit, Cat# 12162).

### Establishment of *EGLN1* KO line of human cells

Human embryonic kidney 293 cells containing the SV40 large T antigen cells (HEK-293T) (ATCC, Cat# CRL-3216) (p27) were seeded at 2x10^4^ cells/cm^2^ in a 12 well plate and cultured with fresh Dulbecco’s Modified Eagle Medium (DMEM) (Thermo Fisher Scientific, Cat# 10564029) supplemented with 10% heat inactivated fetal bovine serum (FBS) (GIBCO, Cat# 10082147) (DMEMc) a day before plasmid exposure. On the day of transfection, the media was changed immediately prior to the transfection itself. Each well was transfected with 7.5 μg of one sgRNA plasmid of interest and 7.5 μg of a *Sp*Cas9 encoding plasmid (AddGene plasmid# 48137) through the calcium phosphate precipitation method as previously described (31). After 5 hours of incubation, the media was changed and HEK-293T cells were provided with fresh DMEMc. Media was changed every other day and HEK-293T cells were morphologically monitored daily to ensure health of the cultures. Once the HEK-293T cells reached 90% confluence, the cells were passaged using 0.05% trypsin-EDTA (Thermo Fisher Scientific, Cat# 25300062) and counted to ensure high viability, and reseeded into T75 flasks. HEK-293T cells were passaged one subsequent time to ensure the removal of plasmid structures from cell culture and these p29 cells were frozen with a freezing solution of 80% DMEMc, 10% FBS (GIBCO, Cat# 10082147), and 10% dimethyl sulfoxide (DMSO) (Sigma-Aldrich, Cat# D8418).

### Determination of *EGLN1* KO efficiency

KO efficiency of individual *EGLN1* sgRNAs was determined through Tracking of Indels by DEcomposition (TIDE) analysis. Genomic DNA of cells co-transfected with a plasmid expressing sgRNA and another expressing *Sp*Cas9 was isolated and purified through a commercially available kit (Qiagen, QIAamp DNA Mini Kit Cat# 51304) as per the manufacturer’s protocol. The *EGLN1* region of interest was enriched via PCR (Gensee Scientific Corporation, Apex Taq RED Master Mix Cat# 42-137) using primers designed by hand, as defined in **Table 1**. Each 25 µl reaction was supplemented with 0.5 µl DMSO to prevent secondary structure formation for clean PCR amplicons. The approximately 1500bp amplicons were run through agarose gel electrophoresis to ensure a lack of secondary structure formation within the sample and DNA was purified from agarose gel (Takara Bio, NucleoSpin Gel and PCR Clean-up Cat# 740609.250) as per the manufacturer’s protocol. Amplicons were sequenced at the UC Davis Sequencing Core through Sanger Sequencing using the same primers that generated the amplicons. These sequences were first visually validated for whether they produced a clean signal before being subsequently uploaded to the TIDE web tool, which provides rapid assessment of genome editing experiments for a target locus (32). The alignment window was 100bp long and began at the recommended 142bp from the start of the amplicon to ensure signal alignment between the test sgRNA sequence and the mock transfected negative control sequence was established.

**Table 1.**
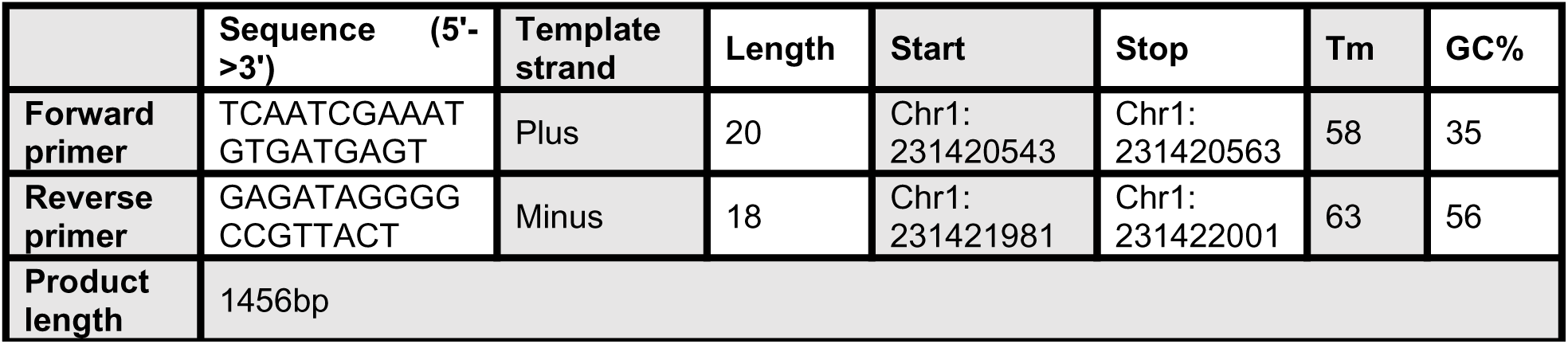
*EGLN1* sgRNA TIDE analysis primer sequences and characteristics

### Identification of sgRNA specific off-targets and quantification thereof

The online web tool Cas-OFFinder was used to identify regions in the human genome that could be loci of off-target double stranded genomic DNA cleavage events (33,34). For each of the three CHOPCHOP identified sgRNA sequences, Cas-OFFinder was used to identify regions where off-targets may occur by specifying *Sp*Cas9 as the gene editor of interest, the human genome being the target genome, and specifying mismatch, DNA bulge and RNA bulge metrics as being 2, 0, and 1 respectively. 6 off-targets were selected for each sgRNA for further investigation and optimized primers were designed for this region through the NIH Primer Design Tool (35,36). KO efficiency at these off-target sites was determined in the same manner described above.

### Per-cell intracellular HIF-1α and secreted protein quantification

Unmodified HEK-293T cells or HEK-293T cells modified by sgRNAs 1, 2, or 3 (Sg1, Sg2, and Sg3 respectively) established in the manner described above were cultured in a T75 flask until 80% confluent and, for each line 2, 6 well plates were plated at passage 30 at a density 20,000 cells/cm^2^. The day following plating, one 6 well plate of each HEK-293T cell line was administered either DMEMc or DMEMc containing 100µM of the chemical hypoxia inducer cobalt chloride (Sigma-Aldrich, Cat# 232696). Two days following the application of these conditional medias, media was removed and frozen for secretomic screening assays for both VEGF-A (R&D Systems, Human VEGF DuoSet ELISA, Cat# DY293B) and VEGF-C (R&D Systems, Human VEGF-C DuoSet ELISA, Cat# DY752B), as per the manufacturer’s protocols. Cells were passaged and counted prior to lysis according to the HIF-1α ELISA (R&D Systems, the Human/Mouse Total HIF-1 alpha/HIF1α DuoSet IC ELISA Cat# DYC1935E) manufacturer’s protocol and total intracellular levels of HIF-1α quantified. Total protein level as determined through these ELISAs was then divided by cell count to provide a per-cell quantification of intracellular and secreted protein levels.

### Relative gene expression analysis by quantitative polymerase chain reaction (qPCR)

Unmodified cells or cells modified by Sg1, Sg2, or Sg3 established in the manner described above were expanded in a T75 flask until 80% confluent before being passaged and seeded into 2, 6 well plates at passage 30 at a density of 20,000 cells/cm^2^. The day following plating, one 6 well plate of each HEK-293T cell line was cultured with either DMEMc or DMEMc containing 100µM of the chemical hypoxia inducer cobalt chloride. Two days following the application of these conditional medias, RNA was isolated via commercially available kit (Qiagen, RNeasy Mini Kit Cat# 74104) by lysing the cells in the vessel as per the manufacturer’s protocol. cDNA was synthesized using high-capacity cDNA reverse transcription kit (Thermo Fisher Scientific, Cat# 4368814) and qPCR was conducted using Fast SYBR Green Master Mix (Thermo Fisher Scientific, Cat# 4385610) in an Applied Biosystems QuantStudio5 real-time PCR machine. Primers for the human housekeeping gene 18S and human proangiogenic and tissue homeostatic factors HIF-1α, VEGF-A, VEGF-C, basic fibroblast growth factor (FGF2), matrix metalloproteinase-2 (MMP2), angiopoietin-1 (Ang-1), chemokine receptor type 4 (CXCR4), and stromal cell– derived factor 1 (SDF-1) were used that were either commercially available, published upon previously, or designed by hand (**Table 2**). Gene expression relative to the appropriate negative control (unmodified HEK-293T cells) in either normoxia or hypoxia was calculated via 2^-ΔΔCT^ and each reaction was carried out in duplicate.

**Table 2.**
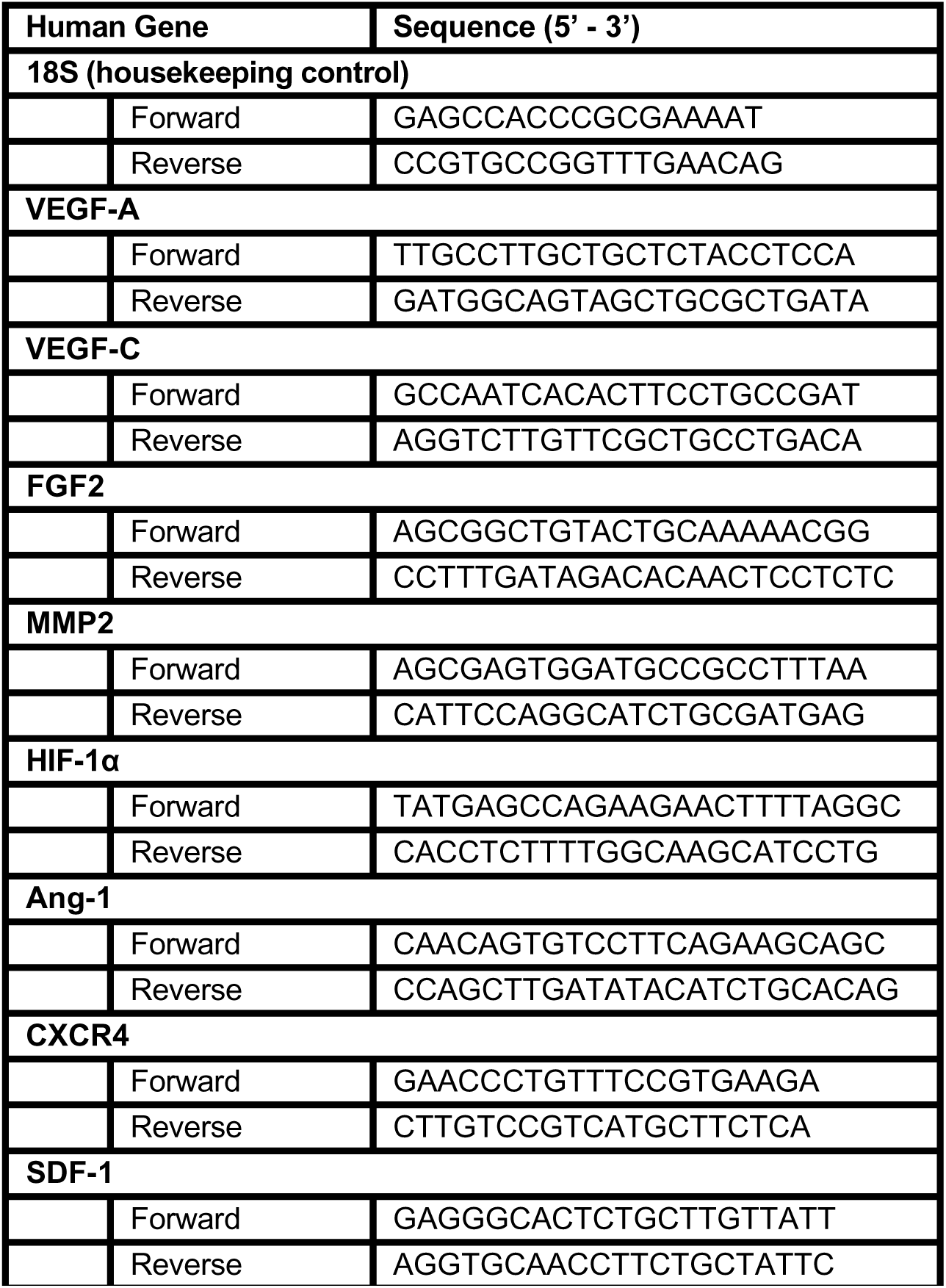
Sequences of the qPCR primers used in this work.

### Estimation of changes in protein expression as a result of *EGLN1* gene editing

Assuming each cell is an isolated system allows for a per-cell quantification estimation of total intracellular or secreted protein changes as a result of gene editing was determined through algebraic estimation as per the following equation:

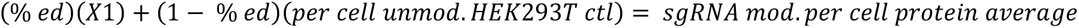

The total protein per-cell average as determined through ELISA is placed on the left-hand side of the equation, and gene editing response is estimated using the *EGLN1* editing efficiency (*%ed*) provided through TIDE analysis. The percent of HEK-293T cells that were unmodified (*unmod.*) were assumed to contain or secrete protein at the same per-cell level of unmodified HEK-293T controls. A simple algebraic rearrangement of the above equation allows for a general estimation of changes in protein levels due to gene editing denoted by *X1*.

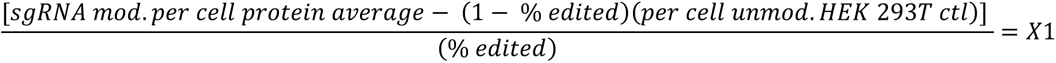

Given that this equation requires the assumption that cellular crosstalk does not impact the internal regulation and secretion of signaling molecules, it should be interpreted as a general estimation.

### Proliferation assay

Proliferation assay as previously described was used to assess the neovascularization potential of *EGLN1* KO HEK-293T cell cultures (37). Adult human dermal microvascular endothelial cells (HMVEC-d) (LONZA, Cat# CC-2543) (passage 8) were cultured until ∼80% confluence before being passaged and seeded at 10,000 cells/cm^2^ in EGM-2MV (LONZA, Cat# CC-3202) in 6-well tissue culture plates and were allowed to adhere for 5 hours before induction of serum starvation in EBM-2 (LONZA, Cat# CC-3156) for 16 hours prior to being administered media of interest, as described below. Media was changed daily for 3 days before cells were detached with 400 μL of 0.05% Trypsin-EDTA (Life Technologies, Cat# 25300054) per well and before samples were collected and counted. The total number of cells in each well was quantified with a Countess automated cell counter (Life Technologies, Cat# AMQAX1000) and averaged per condition (n = 6).

Each of the medias used in this study were either unmodified to test normoxic response or contained 100µM of the chemical hypoxia inducer cobalt chloride. EBM-2 media with FBS and antibiotic (N-media) was preconditioned through incubation with Sg1 modified HEK-293T cells for two days and served as the experimental group. Two negative controls were used – N-media preconditioned through incubation with unmodified HEK-293T cells, and non-preconditioned N-media. EGM-2MV served as a positive control.

### Migratory assay

A modified wound-healing assay was conducted to determine cellular motility in response to media preconditioned as described in the above section. HMVEC-d (passage 9) cells were seeded at 10,000 cells/cm^2^ in 12 well plates with EGM-2MV media, which was changed daily, until 80% confluent. Cells were subsequently starved by incubation with serum-free EBM-2 medium for 16 hours. A scratch was applied manually down the center of each well with a 200 µl micropipette tip in one swift motion as previously described (37). The wells were washed twice with phosphate buffered saline (PBS) (Thermo Fisher Scientific, Cat# 21600010) to remove nonadherent cells. The media of interest was then applied and pictures (10X) were taken using an Axio Vert. A1flourescent microscope (Zeiss, Cat# 491237-0001-000) at two locations along the scratch wound immediately after scratching. The cells were incubated with the medias described above for 24h before being washed twice with PBS and then fixed overnight in 4% formaldehyde (Sigma-Aldrich, Cat# 100496) at 4°C. Cell migratory activity was quantified through pictures taken at the same location through plate bottoms marked with grids (4 mm X 8 mm) – allowing pictures to be taken at the same spot at each time-point for cellular movement.

### Statistics

All results are shown as means with standard deviations. Comparisons between groups were assessed through Student’s unpaired t-tests. All analyses were performed using GraphPad Prism software (GraphPad Software Inc., Version 9.1.2) and differences between conditions are significant if p<0.05.

## RESULTS

### *In silico* generation of human *EGLN1* sgRNA and production thereof

The top three human *EGLN1* CRISPR/Cas9 target site candidates were identified using CHOPCHOP web tool. These candidates and the site-directed mutagenesis primer sequences used to introduce the target sequence to the lentiviral expression cassettes, are described in **Table 3**. The three candidates were selected because they lack off-target cleavage events and have relatively high on-target cleavage efficiencies. The guides (sgRNAs) generated by CHOPCHOP were designed through KO mode, resulting in guides designed to induce a frameshift mutation that renders the protein encoded by this gene non-functional.

**Table 3.**
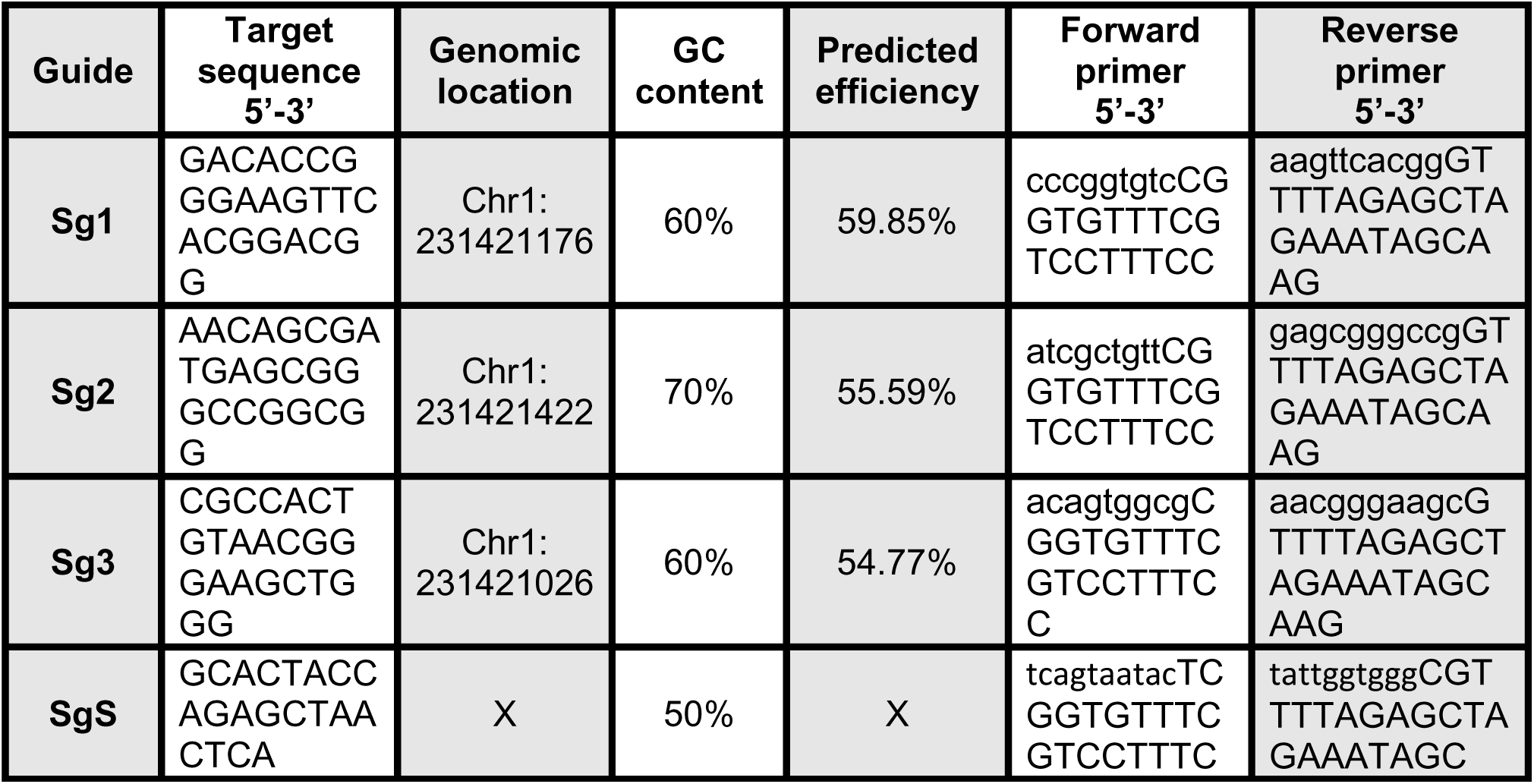
*EGLN1* sgRNA sequences and scramble (“SgS”) sequence characteristics and mutagenesis primer sequences. Lowercase letters are the insertion region, uppercase is the sequence specifying the insertion region into the plasmid backbone generated *in silico*.

### Genomic modification capabilities of *EGLN1* sgRNAs

All three sgRNA candidates were tested for their maximal knockout capabilities by transfection in HEK-293T cells. HEK-293T cells are relatively easy to transfect and have previously been utilized in studying human *EGLN1* (38–40). TIDE analysis assessed on-target genomic modification capabilities of all *EGLN1* sgRNAs.

TIDE analysis determined that all three sgRNAs introduced on-target indel mutations in the single digits. Sg1 had a total efficiency of 3.4% with a significant occurrence of a single base pair insertion. Sg2 displayed a total efficiency of 3.9% with no singular statistically significant indel. Finally, Sg3 showed a total efficiency of 5.1% with a significant occurrence of a single base pair deletion (**Figure 2**).

**Figure 2.**
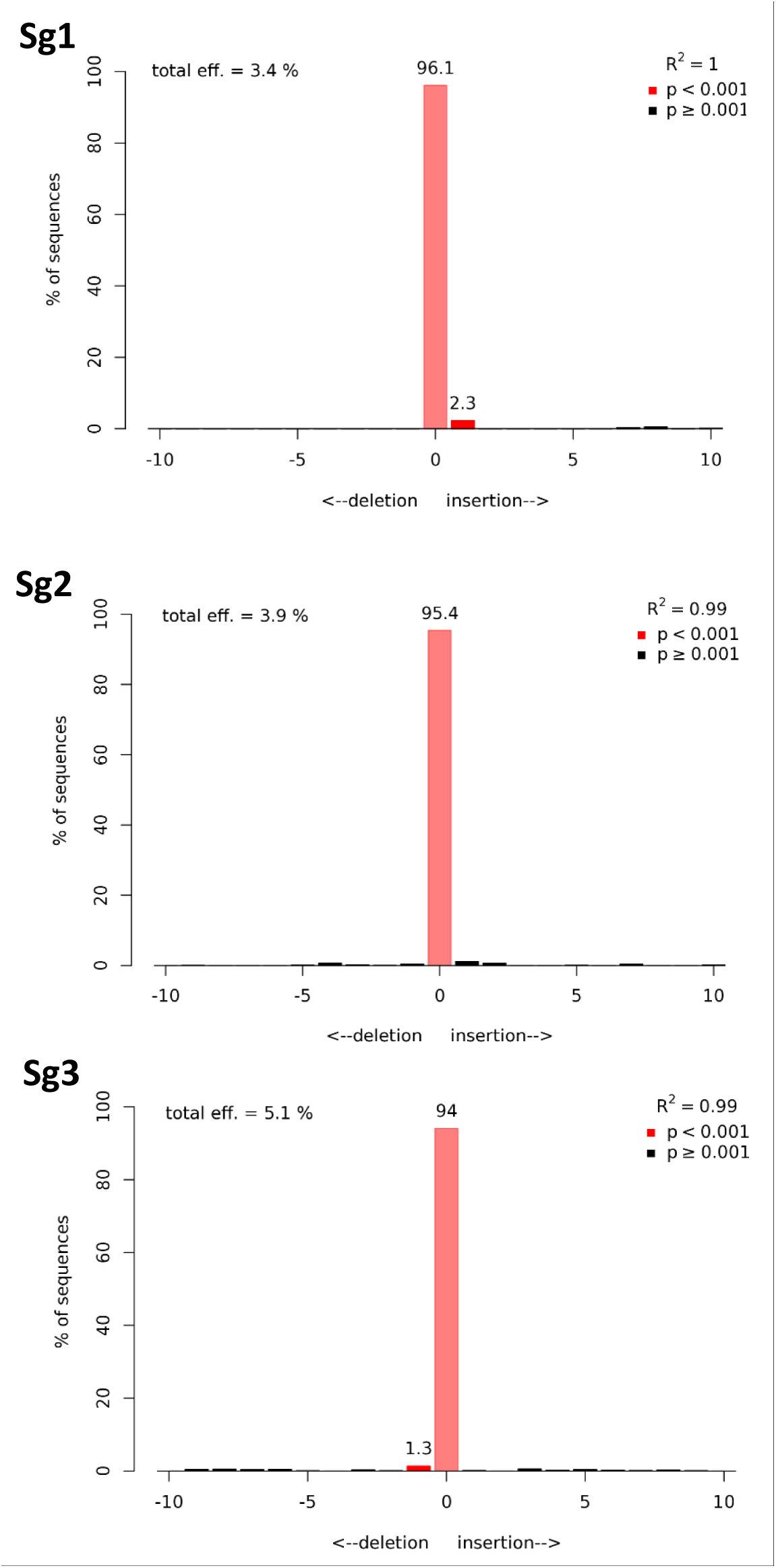
TIDE analysis of on-target *EGLN1* KO of the three *in silico* generated sgRNAs (Sg1, Sg2, and Sg3) in HEK-293T cells.

Cas-OFFinder generated six potential off-targets per guide under 2 potential base pair mismatches between the sgRNA and genomic region and 1 RNA base pair bulge. A summary of indel rates, as determined through TIDE analysis of all on-target and successful off-targets primer sets, is presented in **Table 4**. Most predicted off-targets for the three guides presented had low occurrence of indel mutations. However, Sg1 predicted off-target number 5 showed an indel rate of 9%, Sg2 predicted off-target 5 showed an indel rate of 8.6%, and Sg3 predicted off-target numbers 5 and 6 showed an indel rate of 15.5% and 8.2%, respectively (**Figure 3**).

**Figure 3.**
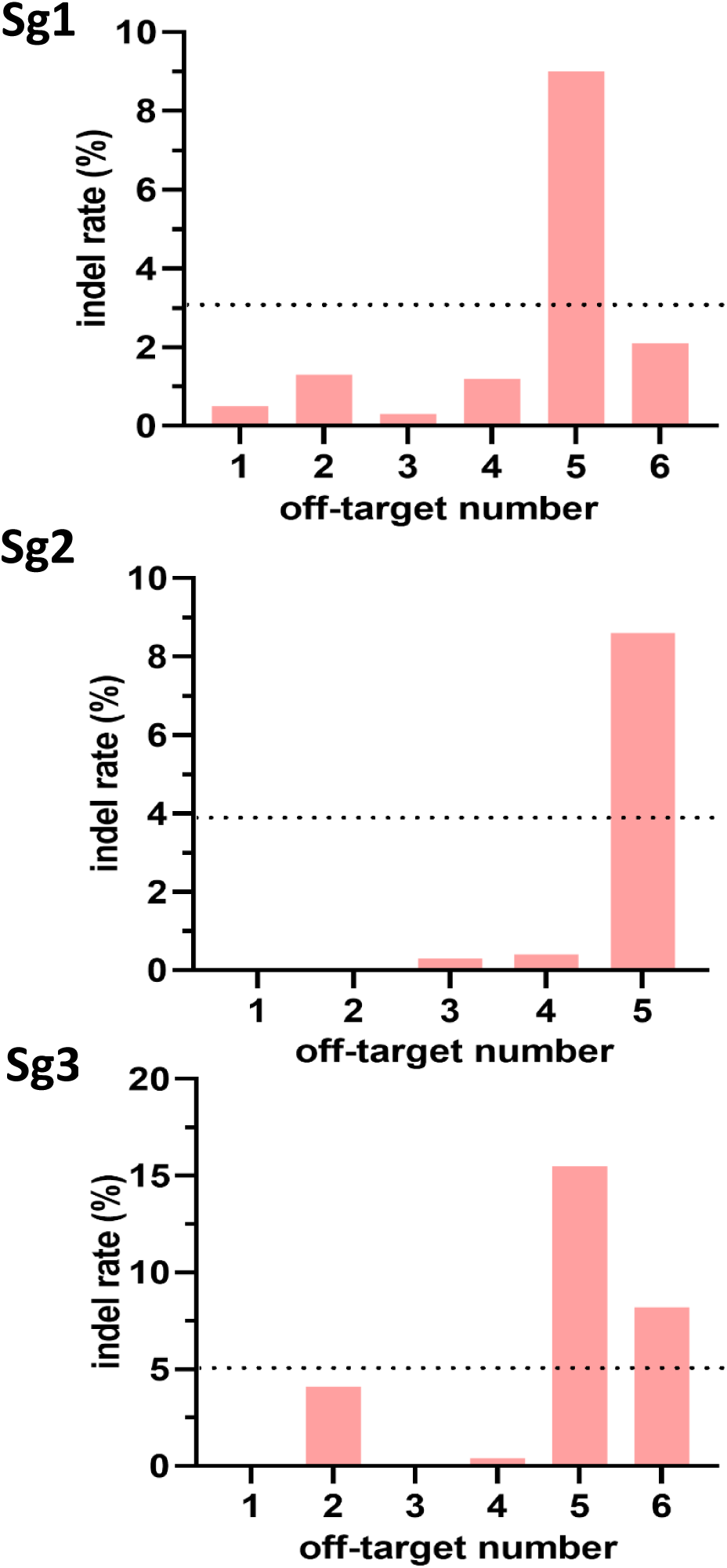
TIDE analysis results of off-target indel rates of Sg1, Sg2, and Sg3 in HEK-293T cells. Bars represent indel rate per Cas-OFFinder predicted off-target site, dashed lines represent respective on-target indel rate for reference.

**Table 4.** Prevalence of Cas-OFFinder predicted off-target cleavage events as determined by TIDE analysis.

| Guide | On-target | Off-targets |
| --- | --- | --- |
| **Sg1** | 3.4% | 0.5%, 1.3%, 0.3%, 0%, 9%, 2.1% |
| **Sg2** | 3.9% | 0%, 0%, 0.3%, 0.4%, 8.6% |
| **Sg3** | 5.1% | 0%, 4.1%, 0%, 0.4%, 15.5%, 8.2% |

### Biomolecular regulatory changes resulting from *EGLN1* KO

Biomolecular regulatory modifications were examined in HEK-293T cells due to gene editing by *Sp*Cas9 localized to *EGLN1* by the proposed sgRNA candidates.

Proliferation of all respective lines of HEK-293T cells established in the manner described above was determined (**Figure 4**) for per-cell protein-level quantification. Only Sg2 stimulated a significant proliferative response in normoxia compared to unmodified controls in both normoxic and hypoxic conditions, however, this significance was not maintained in hypoxia. No statistical significance was seen between any group in normoxia and hypoxia.

**Figure 4.**
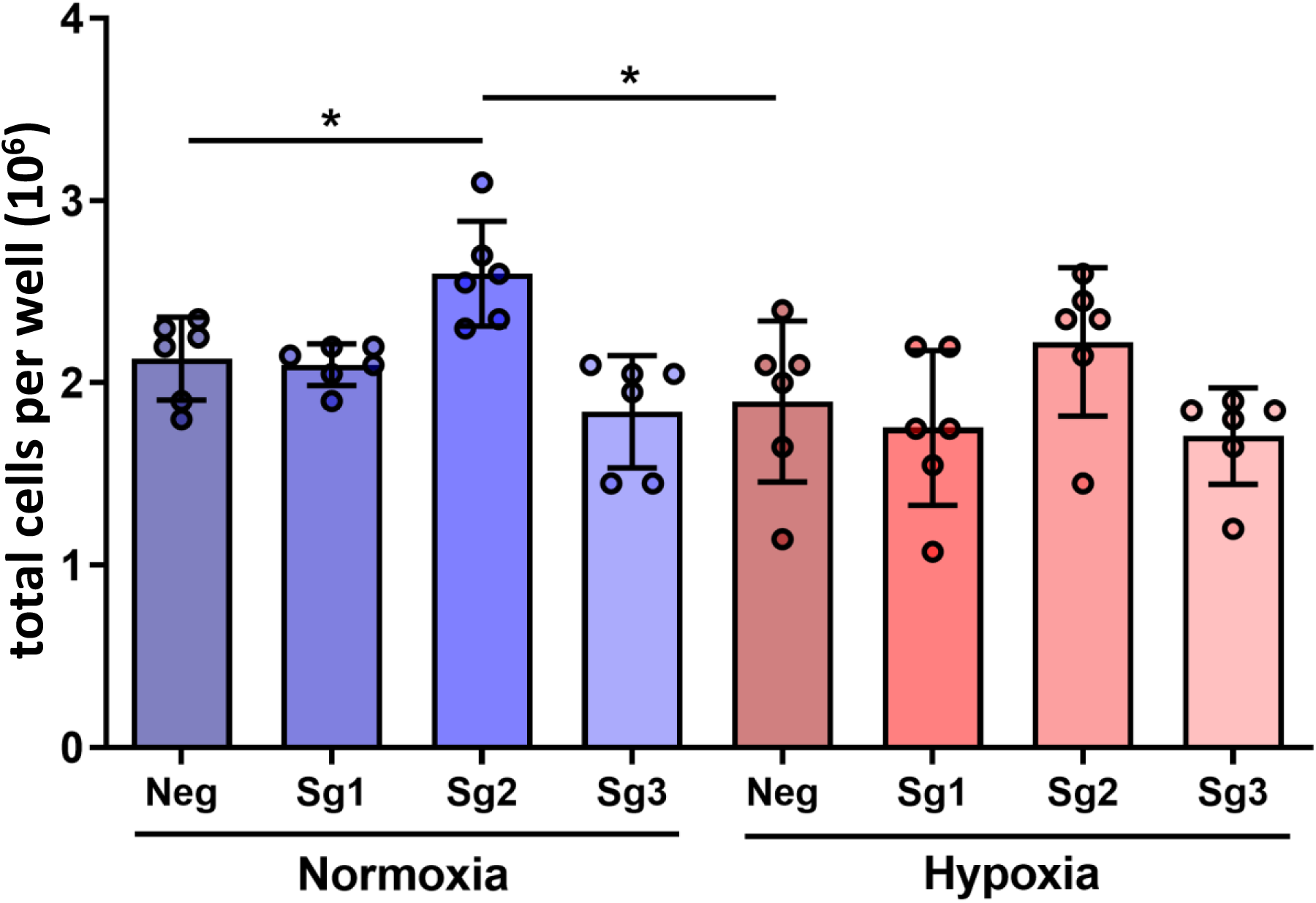
Total number of either unmodified (Neg) or guide modified (Sg1, Sg2, and Sg3) HEK-293T cells per well of a 6 well plate from which cell lysates and media were isolated for protein quantification assays. Columns represent mean, points represent individual wells (n=6), bars represent standard deviation. * represents significance at p<0.05.

Given that *EGLN1* encodes the core regulator of HIF-1α and that upregulation of HIF-1α was the preeminent goal of this system, quantification of HIF-1α upregulatory capabilities of all guide sequences was the primary determinant of functional KO success. Per-cell intracellular HIF-1α protein levels (**Figure 5**) as determined by ELISA show that Sg1 and Sg3 could stimulate a significant upregulation of the HIF-1α target protein in normoxia when compared to the unmodified control, and this upregulation was maintained in hypoxic conditions. Importantly, the upregulation of HIF-1α Sg1 and Sg3 introduced in normoxic conditions was not significantly different from the endogenous hypoxic HIF-1α upregulatory response of unmodified controls.

**Figure 5.**
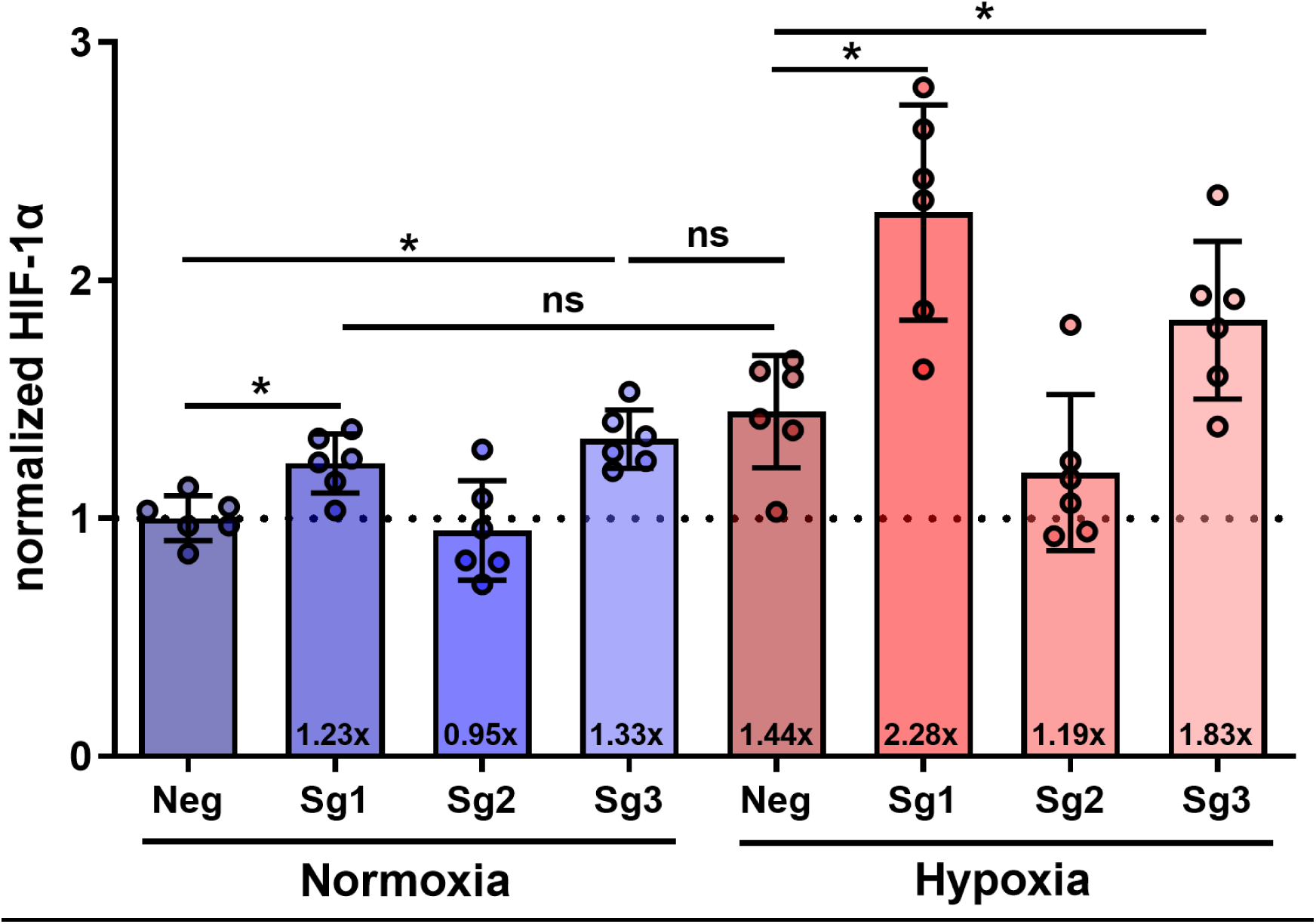
*EGLN1* KO (Sg1, Sg2, or Sg3) resulted in increased intracellular levels of HIF-1α in HEK-293T cells in the cases of two of three guides when normalized to the per-cell, normoxic, unmodified HEK-293T negative control (Neg) as quantified by ELISA for the proangiogenic master regulatory protein HIF-1α. Columns represent mean, points represent individual wells (n=6), bars represent standard deviation. * represents significance at p<0.05.

To investigate the impact of HIF-1α upregulation on downstream regulatory activities in response to *EGLN1* KO, media of the above cell cultures was collected and screened to quantify differences in secreted factors. Cells modified by Sg1 and Sg3 secreted significantly more of the core proangiogenic protein VEGF-A compared to the unmodified control in normoxia, and Sg1 maintained a significant increase in secretion compared to the unmodified control in hypoxia (**Figure 6**). Slight upregulatory trends were also seen in VEGF-C protein secretion in Sg1 and Sg3 modified groups compared to the unmodified control in normoxia, which was maintained in hypoxia for Sg1 (**Figure 7**). However, no changes were significant between any group, including the normoxic and hypoxic responses of the unmodified control group.

**Figure 6.**
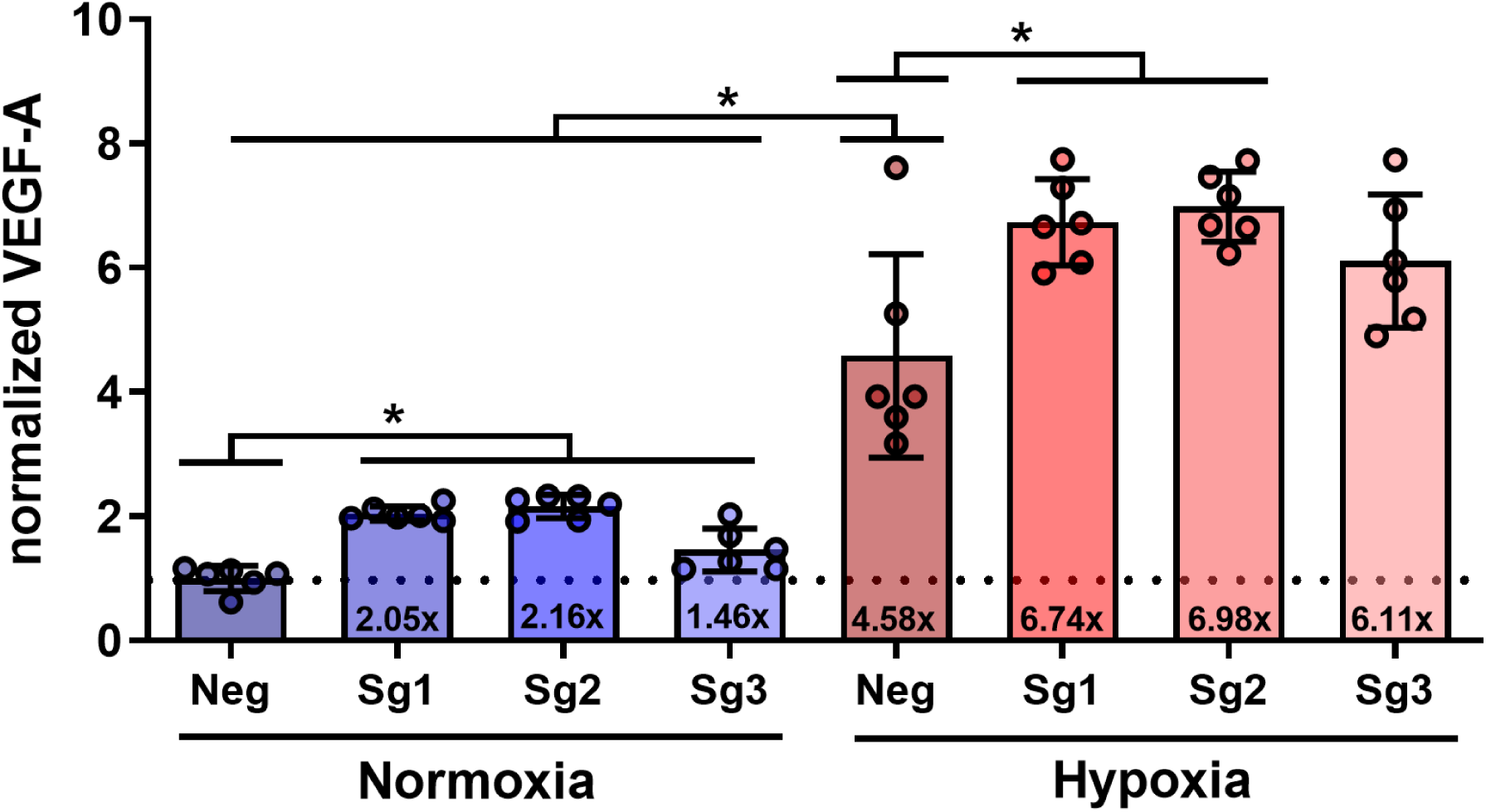
*EGLN1* KO (Sg1, Sg2, or Sg3) increased VEGF-A secretion in HEK-293T cells for all guides in normoxic conditions and for two of three guides in hypoxic conditions when normalized to the per-cell, normoxic, unmodified HEK-293T negative control (Neg), as quantified by ELISA for the strongest-identified-to-date proangiogenic protein VEGF-A. Columns represent mean, points represent individual wells (n=6), bars represent standard deviation. * represents significance at p<0.05.

**Figure 7.**
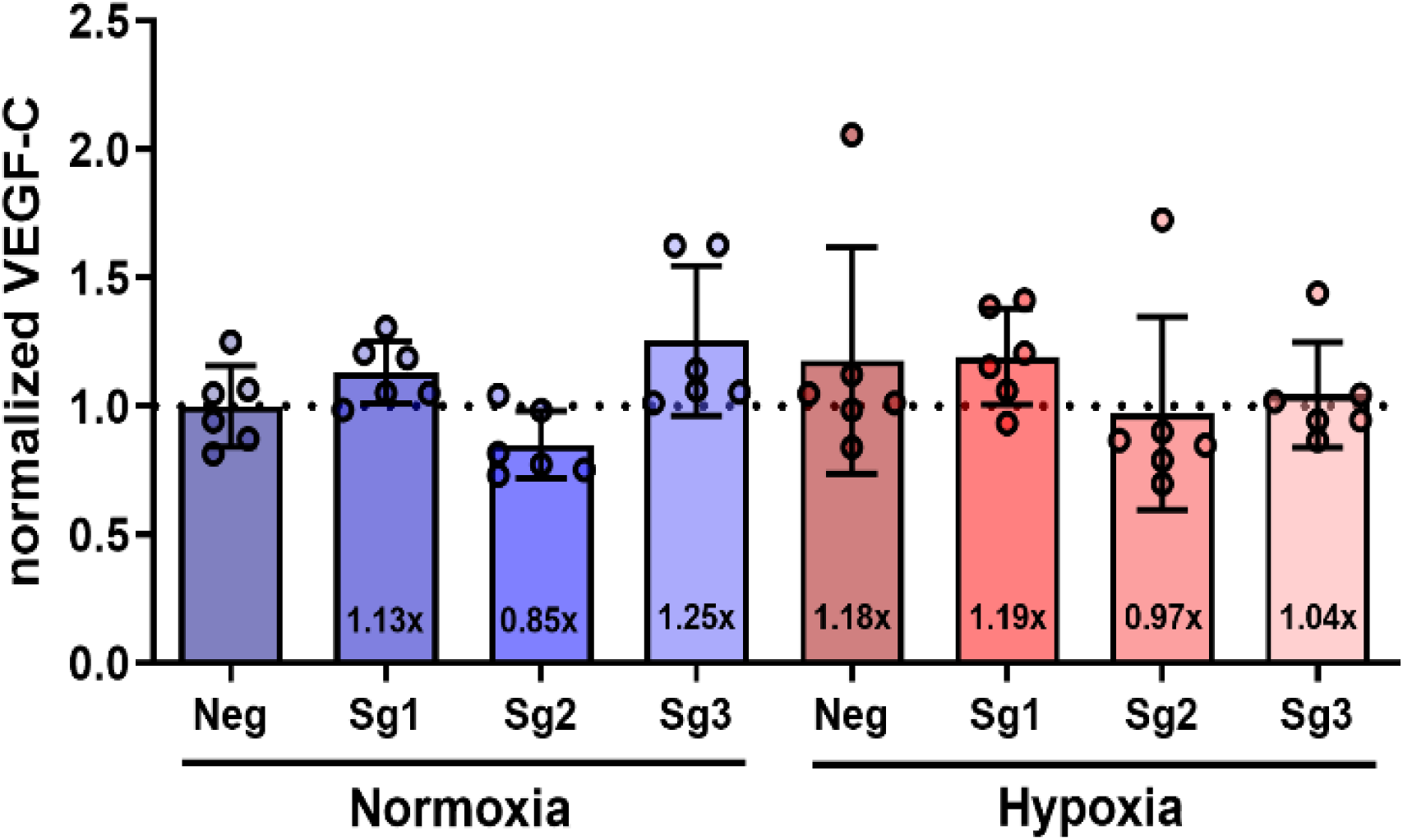
VEGF-C secretion response was unchanged in HEK-293T cells in response to *EGLN1* KO (Sg1, Sg2, or Sg3) when normalized to the per-cell, normoxic, unmodified HEK-293T negative control (Neg) as quantified by ELISA for the tissue homeostatic protein VEGF-C. Columns represent mean, points represent individual wells (n=6), bars represent standard deviation.

To investigate the downstream transcriptional upregulation potential of *EGLN1* KO, the mRNA of eight HIF-1α associated proangiogenic and homeostatic factors was quantified through qPCR in both normoxic and hypoxic conditions (**Figure 8**). Upregulation in normoxic conditions was seen in seven of the eight factors, four of which, VEGF-A, FGF2, MMP2, and CXCR4, were statistically significant compared to the unmodified negative control. Taking Sg1 editing efficiency into consideration shows that *EGLN1* KO cells are estimated to contain 10.4x more VEGF-A, 3.4x more VEGF-C, 14.5x more FGF2, 17.1x more MMP2, 5.1x more HIF-1α, 1.4x less Ang-1, 17.7x more CXCR4, and 2.8x more SDF-1 transcripts in normoxic oxygen tensions. In hypoxic conditions, transcript levels of all significantly upregulated factors in normoxia were not significantly different from the unmodified negative control, and three factors – VEGF-C, HIF-1α, and Ang-1 – were significantly downregulated. Taking Sg1 editing efficiency into consideration, *EGLN1* KO cells are estimated to contain 2.2x less VEGF-A, 4.6x less VEGF-C, 1.9x more FGF2, 0.1x more MMP2, 4.3x less HIF-1α, 7.5x less Ang-1, 1.6x less CXCR4, and 2.5x less SDF-1 transcripts compared to unmodified controls in hypoxic oxygen tensions.

**Figure 8.**
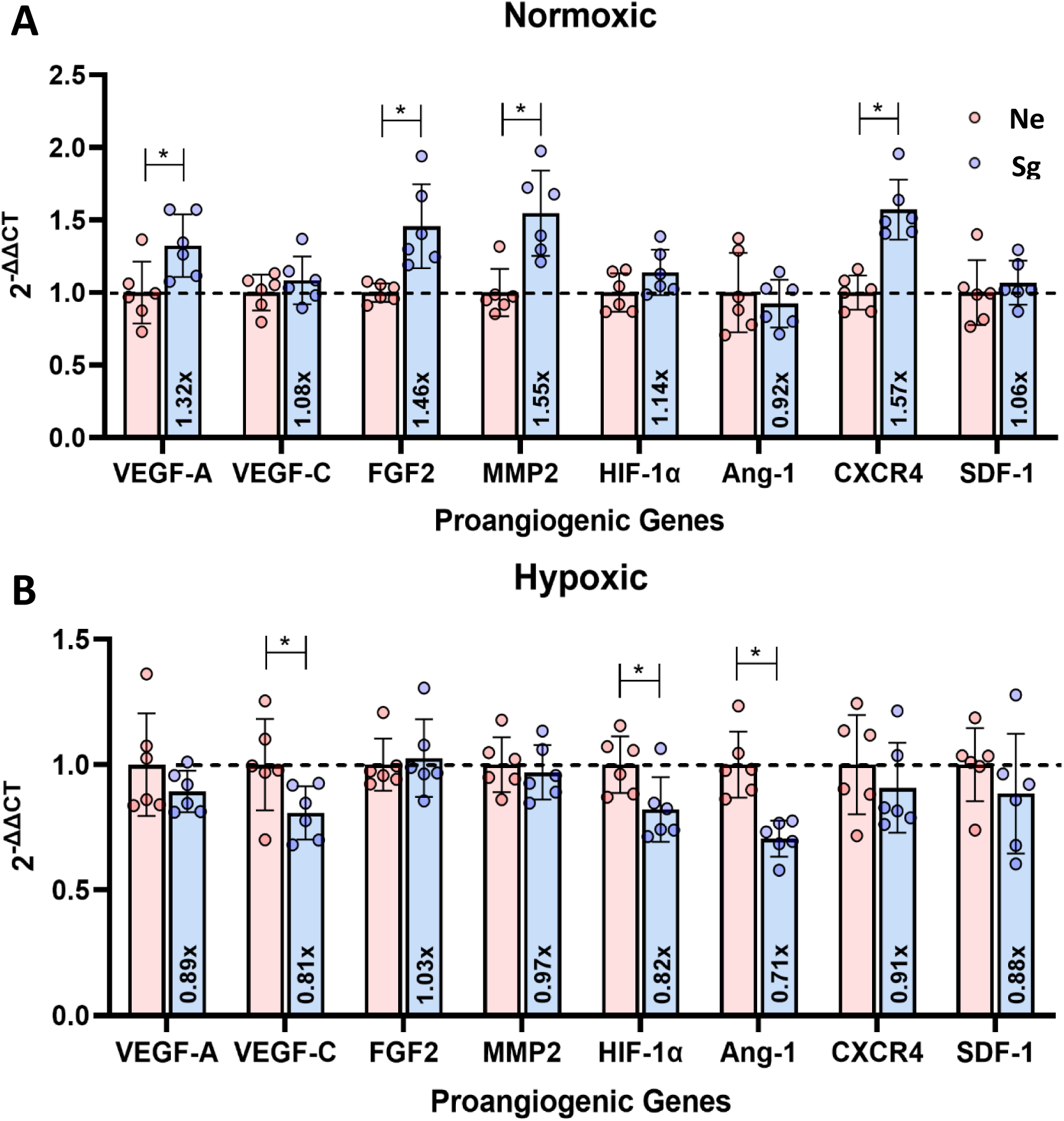
Changes in HEK-293T mRNA levels of select proangiogenic, tissue homeostatic, ECM remodeling, and immunomodulatory factors in response to *EGLN1* KO by Sg1 compared to unmodified HEK-293T (Neg) in normoxia (A) and hypoxia (B). Columns represent mean, points represent individual wells (n=6), bars represent standard deviation. * represents significance at p<0.05.

#### Proliferative response

To investigate the functional effect of *EGLN1* KO secretomic modifications, we feed the preconditioned media from *EGLN1* KO HEK-293T cells to HMVEC-d cells cultured under normoxic or chemically hypoxic conditions, while using preconditioned media from unmodified HEK-293T cells as a control. Preconditioned media from the Sg1 *EGLN1* KO HEK-293T cells significantly enhanced HMVEC-d proliferation in normoxic oxygen tensions compared to unmodified cells (**Figure 9**). Interestingly, this HMVEC-d proliferative response was not significantly different from preconditioned media of Sg1 *EGLN1* KO HEK-293T cultured in hypoxic conditions, nor from preconditioned media of unmodified HEK-293T cells.

**Figure 9.**
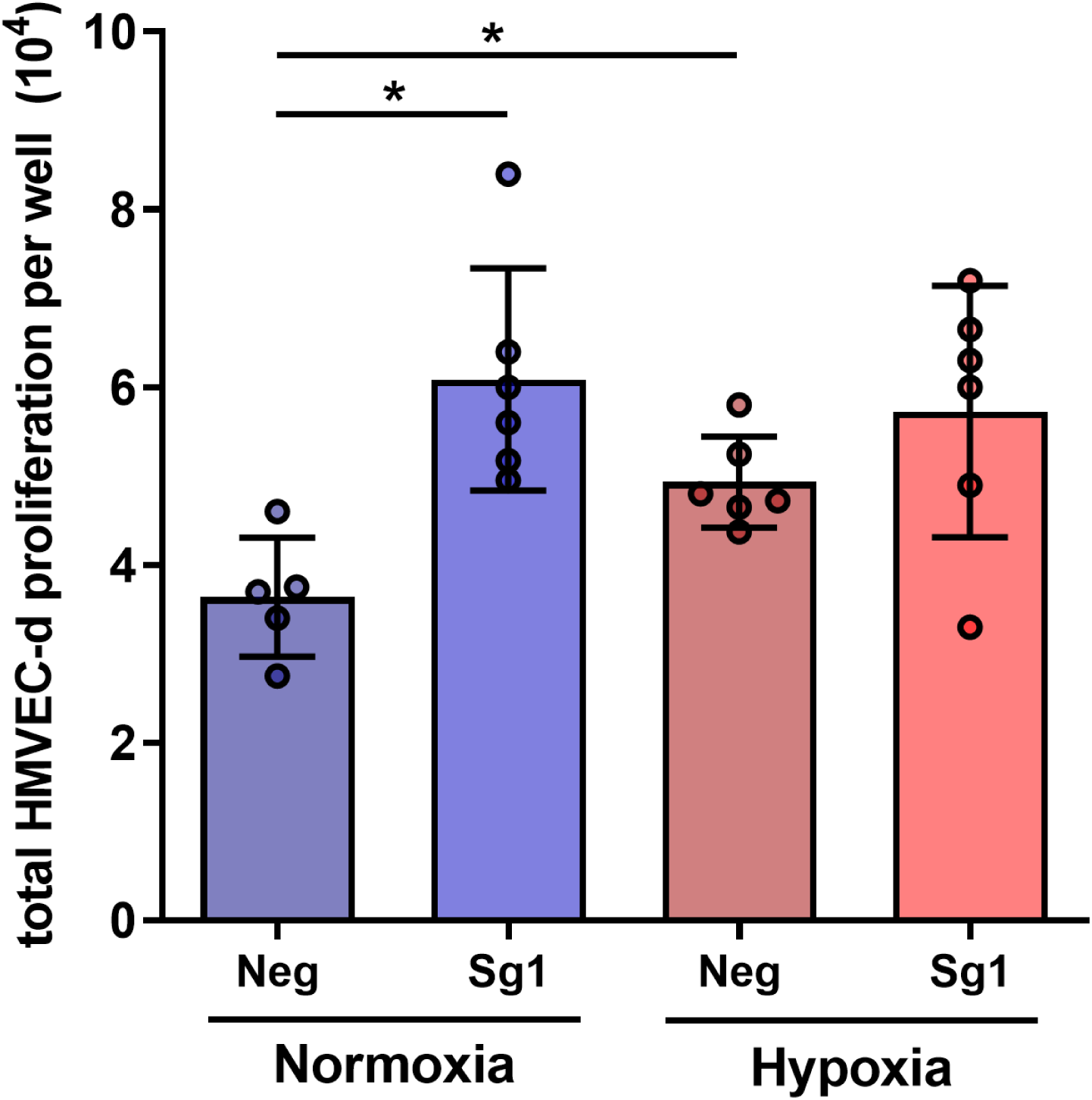
HMVEC-d proliferation response was greater when exposed to *EGLN1* KO (Sg1) HEK-293T preconditioned media compared to exposure to unmodified (Neg) HEK-293T preconditioned media in both normoxic and hypoxic conditions. Columns represent mean, points represent individual wells (n=6), bars represent standard deviation. * represents significance at p<0.05.

#### Migratory activity

In both normoxic and hypoxic oxygen tensions, preconditioned media from *EGLN1* KO HEK-293T cells increased motility trends in a migratory assay of HMVEC-d cells compared to preconditioned media from unmodified HEK-293T negative control cells (**Figure 10**). No statistically significant differences were seen between any groups in any condition.

**Figure 10.**
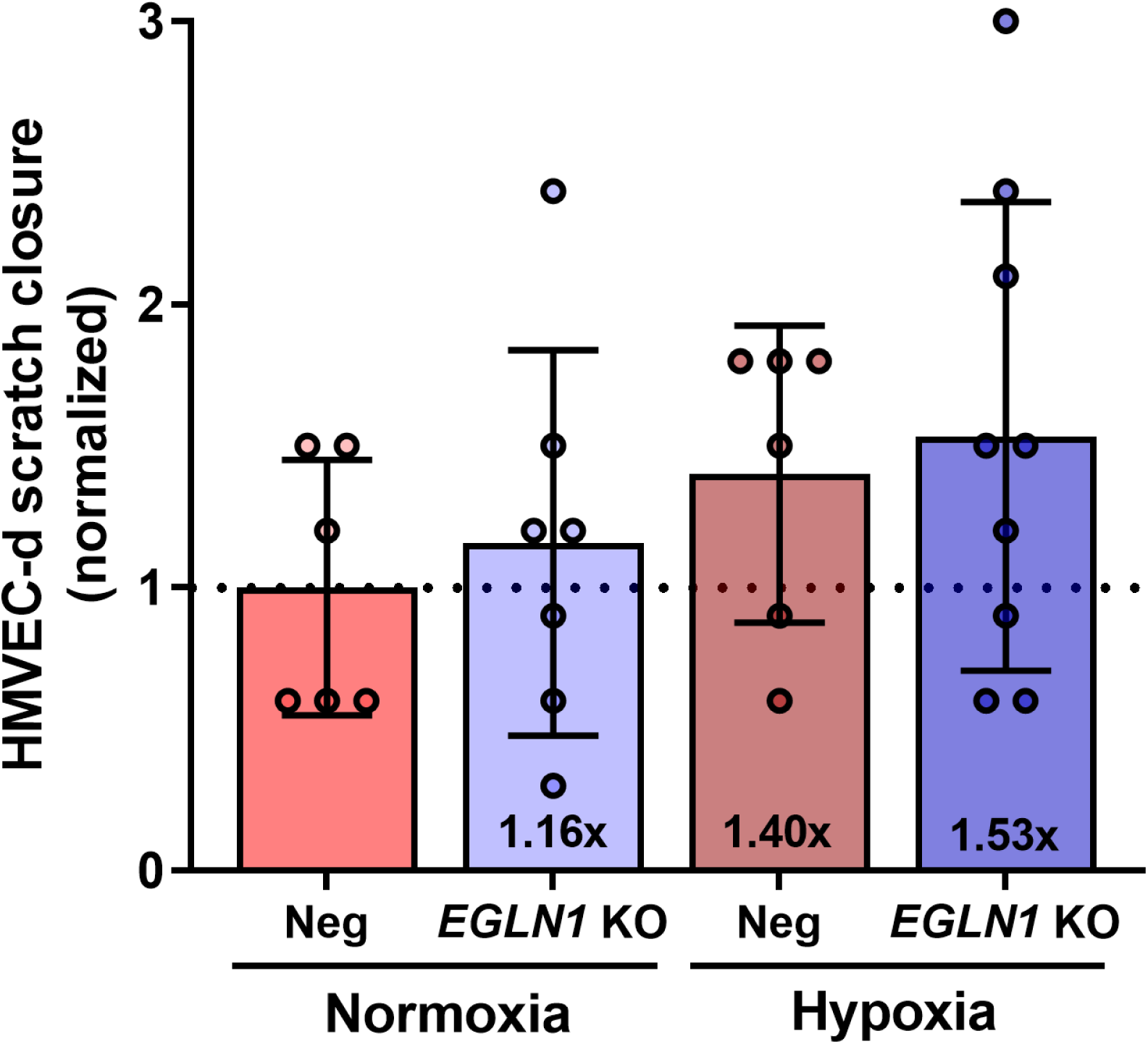
HMVEC-d scratch closure was greater when exposed to preconditioned media from *EGLN1* KO (Sg1) HEK-293T cells versus unmodified (Neg) HEK-293T cells in both normoxic or hypoxic conditions. Columns represent mean, points represent individual image points (n=6-9), bars represent standard deviation.

## DISCUSSION

The results of this manuscript suggest that even marginal disruptions to the HIF-1α pathway at the genomic level of *EGLN1* can initiate a significant proangiogenic response. To our knowledge, this study is the first to investigate *EGLN1* KO to determine the likelihood of a permanent intervention of this pathway as an *in situ* regenerative therapeutic option. Specifically, this study demonstrates the potential of *EGLN1* gene editing using *Sp*Cas9 to induce intracellular HIF-1α protein levels and pro-angiogenic protein secretion, highlighting the impact of such proteins on human endothelial cell proliferation and migration. Our findings support that *EGLN1* is a key regulator of angiogenesis and that its inhibition could be a viable therapeutic strategy for various diseases characterized by impaired vascular function.

In our proof-of-concept analysis, the three *in silico* designed *EGLN1* sgRNA constructs achieved on-target editing rates between 3 – 6% as determined through TIDE analysis. Each guide had a predicted off-target genomic location with a higher editing rate than each on-target editing efficiency. However, if disturbed, no off-target activity occurred in protein coding regions known to cause pathogenesis in ischemic tissues. We acknowledged the intrinsic limitations of using a small number of potential *EGLN1* sgRNAs and understand that some may argue this restricts the scope of this work. However, this foundational work is critical for validating further studies.

This study demonstrates that, in the context of tissue ischemia where low oxygen tensions are clinically relevant, human cellular response to *EGLN1* KO by Sg1 can significantly upregulate HIF-1α in both normoxic and hypoxic oxygen tensions when compared to unmodified controls in those same conditions. Importantly, the upregulatory response induced by *EGLN1* KO was not significantly different from endogenous hypoxic HIF-1α upregulatory response, suggesting that *EGLN1* KO can synthetically induce an intracellular hypoxic state. Considering the 3.4% Sg1 editing efficiency, our calculations indicate that, in normoxic conditions, Sg1 edited cells contain an average of 7.8x more HIF-1α than non-edited cells. This level increases to 18.2x in hypoxic conditions, demonstrating strong HIF-1α upregulation potential. This upregulatory trend is consistent with what has been previously documented in conventional HIF-1α integrative gene therapy strategies (41).

Our findings demonstrate that *EGLN1* KO by Sg1 elicits a strong proangiogenic protein secretory response. On a per-cell basis, Sg1-modified cells secrete twice the amount of VEGF-A protein compared to unmodified controls in normoxic conditions.

Considering a 3.4% editing efficiency, Sg1-edited cells secrete 31.9x more VEGF-A than their unmodified counterparts in normoxia, and 14.9x more in hypoxia. Given that VEGF-A is the most potent proangiogenic protein described to date, these results demonstrate strong proangiogenic potential (42). Modifications directed at the master regulatory level show promise for therapeutic angiogenesis applications that could be stronger than traditional gene therapy interventions. For example, in human mesenchymal stromal cells (MSCs), cells transduced with lentivectors encoding VEGF-A secreted approximately 10x more VEGF-A than unmodified MSCs (43). Bone marrow stem cells (BMSCs) transduced at 90% efficiency with integrating lentivectors encoding HIF-1α displayed a 7x increase in VEGF-A protein secretion (44). Similarly, in another study, BMSCs transduced at 80% efficiency to express HIF-1α stimulated 6x more VEGF-A secretion (41). Therefore, both conventional VEGF-A and HIF-1α gene therapy methods do not stimulate secretion responses as strongly as *EGLN1* KO. These findings suggest that *EGLN1* KO warrants further investigation to upregulate proangiogenic factors significantly.

Tissue homeostatic protein VEGF-C expression was also altered in response to *EGLN1* KO, though not significant on a per-cell level. Considering the 3.4% Sg1 editing rate, edited cells secrete 4.8x more VEGF-C than unmodified cells in normoxia, and 1.5x more in hypoxia. These differences in VEGF-C secretion could have potential biological implications given that VEGF-C plays a pivotal role in regulating lymphatic vessel growth and mediating interstitial pressure, which, if left uncontrolled, can hamper therapeutic agent delivery and distribution in affected tissues (45,46). Therefore, upregulation of VEGF-C and other homeostatic proteins impacted by the HIF-1α pathway are essential for a coordinated tissue-scale response necessary for therapeutic angiogenesis.

*EGLN1* KO cells secreted factors that enhanced neovascularization activity in primary human endothelial cells. Normoxic proliferation in HMVEC-d cells incubated with preconditioned media from *EGLN1* KO cells was significantly increased compared to those incubated with preconditioned media from unmodified HEK-293T cells. Interestingly, this proliferative effect seen in HMVEC-d groups cultured with media from normoxic *EGLN1* KO preconditioned media was not significantly different from HMVEC-d cell response to HMVEC-d groups cultured with preconditioned media from unmodified HEK-293T cells, nor hypoxic *EGLN1* KO cells. Although there was a statistically significant difference in proliferation between normoxic and hypoxic groups of HMVEC-d cells cultured in preconditioned unmodified HEK-293T cells, there was no difference between HMVEC-d groups receiving *EGLN1* KO media. These results are promising because they demonstrate that even small editing efficiencies – such as the 3.4% editing rate described in this work – can stimulate a normoxic secretomic response robust enough to signal endothelial cells in a manner comparable to an endogenous hypoxic response. Migratory assay results also suggest the potential of increased motility upon exposure to secreted factors from *EGLN1* KO cells. *EGLN1* KO preconditioned media groups demonstrate higher motility trends than their unmodified counterparts in both normoxia and hypoxia. However, to fully capture the potential motility aspect of this work, further research should expand these initial proof-of-concept analyses.

In normoxia, upregulation of mRNA levels correlated to upregulation of protein expression of HIF-1α (5.19x more mRNA and 8x more protein), VEGF-A (11.3x more mRNA and 35x more protein), and VEGF-C (3.6x more mRNA and 5x more protein), which is consistent with data of conventional gene therapies for HIF-1α in normal physiological oxygen tensions (15,41,44,47). However, in hypoxia, mRNA levels compared to unmodified controls are inversely correlated for all three proteins, which is somewhat contrary to previous reports of elevated HIF-1α and proangiogenic factor mRNA and protein levels in response to HIF-1α gene therapy in hypoxic tissues (48). The discrepancies between our findings and the previous study could be due to methodological differences related to unique strategy used in our study.

The interdependence of multiple factors in the HIF-1α pathway could be responsible for the observed discrepancies between mRNA and protein levels in hypoxic and normoxic conditions. The regulatory feedback loop involving HIF-1α and *EGLN1* might also contribute to some extent to the observed differences (16). As such, permanent modifications within this pathway can possibly produce expression profiles unique to both normoxic and hypoxic responses. Though studies on global and conditional knockouts of this gene are well tolerated in animal models and demonstrate hypervascularity potential (21,22,49), detailed quantitative analyses of these transgenics’ mRNA and protein expression profiles in normoxic and hypoxic conditions are lacking. Thus, to better understand the unique responses of *EGLN1* KO cells, future studies should focus on broader mRNA quantification assays that can elucidate the complex regulatory mechanisms involving the HIF-1α pathway.

## CONCLUSION

This study successfully demonstrated that *EGLN1* KO by *Sp*Cas9 can stimulate HIF-1α mediated regulatory activity in human cells at both oligonucleotide and protein levels. The robust hypoxic response observed even at low editing efficiencies provides a strong proof-of-concept for further investigation into the short and long-term biomolecular regulatory changes resulting from *EGLN1* genomic editing. The promising results presented herein warrant further investigation of prolonged HIF-1α stabilization methods achieved through permanent modification to *EGLN1* in human cells towards the development of more effective therapeutic angiogenesis strategies.

## CONFLICT OF INTEREST

The authors declare that this manuscript was written in the absence of any commercial or financial relationships that could be construed as a conflict of interest.

## FUNDING

This manuscript was developed with funding from the NIGMS-funded Pharmacology Training Program T32GM099608 (SS) and the American Heart Association Predoctoral grant # 20PRE35210399 (SS). This work was supported by the American Heart Association grant # 19IPLOI34760654 (EAS) and the São Paulo Research Foundation (FAPESP) grant # 2019/10922-9 (RSS).

## Notes

### Competing Interest Statement

The authors have declared no competing interest.

